# Replicable multivariate BWAS with moderate sample sizes

**DOI:** 10.1101/2022.06.22.497072

**Authors:** Tamas Spisak, Ulrike Bingel, Tor Wager

## Abstract

Brain-Wide Association Studies (BWAS) have become a dominant method for linking mind and brain over the past 30 years. Univariate models test tens to hundreds of thousands of brain voxels individually, whereas multivariate models (‘multivariate BWAS’) integrate signals across brain regions into a predictive model. Numerous problems have been raised with univariate BWAS, including lack of power and reliability and an inability to account for pattern-level information embedded in distributed neural circuits^1–3^. Multivariate predictive models address many of these concerns, and offer substantial promise for delivering brain-based measures of behavioral and clinical states and traits^2,3^.

In their recent paper^4^, Marek et al. evaluated the effects of sample size on univariate and multivariate BWAS in three large-scale neuroimaging dataset and came to the general conclusion that *“BWAS reproducibility requires samples with thousands of individuals”*. We applaud their comprehensive analysis, and we agree that (a) large samples are needed when conducting univariate BWAS of individual differences in trait measures, and (b) multivariate BWAS reveal substantially larger effects and are therefore more highly powered. However, we disagree with Marek et al.’s claims that multivariate BWAS provide *“inflated in-sample associations”* that often fail to replicate (i.e., are underpowered), and that multivariate BWAS consequently require thousands of participants when predicting trait-level individual differences. Here we substantiate that (i) with appropriate methodology, the reported in-sample effect size inflation in multivariate BWAS can be entirely eliminated, and (ii) in most cases, multivariate BWAS effects are replicable with substantially smaller sample sizes (Figure 1).

## Main

Marek et al. evaluate in-sample effect size inflation in multivariate BWAS by training various multivariate models in a ‘discovery sample’ and comparing the in-sample effect sizes (prediction-outcome correlation, *r*) to the performance in an independent ‘replication’ sample. Based on a bootstrapping analysis, with variously sized pairs of samples drawn randomly from the Adolescent Brain Cognitive Development study (n=11,874), the authors report a severe effect size inflation of Δr = −0.29 (average out-of-sample minus in-sample r) and conclude that *“even at the largest sample sizes (n ≈ 2,000), multivariate in-sample associations remained inflated on average”*.

The issue with claims of inflation is that Marek et al.’s in-sample effect size estimates were based on training multivariate models on the entire discovery sample, without cross-validation or other internal validation (as confirmed by inspection of the code and discussion with the authors). The problem of overfitting when training multivariate models is well known^5^, and standard practice in machine learning is to evaluate accuracy (and other metrics of fit) on data independent of those used during training^2,5^. Marek et al.’s claim of inflated estimates is thus misleading; the use of widely accepted cross-validation procedures would be expected to reduce average in-sample inflation to zero. Multivariate BWAS on small samples should still produce more variable results^5^ with reduced power, but no systematic bias.

To demonstrate this, and to assess both bias and power in multivariate BWAS, we analyzed functional connectivity data from the Human Connectome Project^6^ (one of the datasets in Marek et al.) with similar analyses to those used by Marek et al., but using cross-validation to estimate in-sample effect sizes. As shown in **Figure 1** (a-d), cross-validated effect size estimates are unbiased (i.e., not inflated on average), irrespective of the sample size and the magnitude of the effect. As expected, smaller sample sizes resulted in lower power (Figure 1e) and increased variability across samples (**Figure 1**c). Such variability is undesirable because it reduces the probability of independent replication (Figure 1f). In addition, selection biases— most notably publication bias—can capitalize on such variability to inflate effect sizes in the literature (**Figure 1**g). Though these effects of using small sample sizes are undesirable, they do not invalidate the use of multivariate BWAS in small samples, and publication biases can be mitigated by practices that are quickly becoming standards in the field^2,5^, including pre-registration, registered reports, and the use of hold-out samples tested only once.

**Figure 1.**
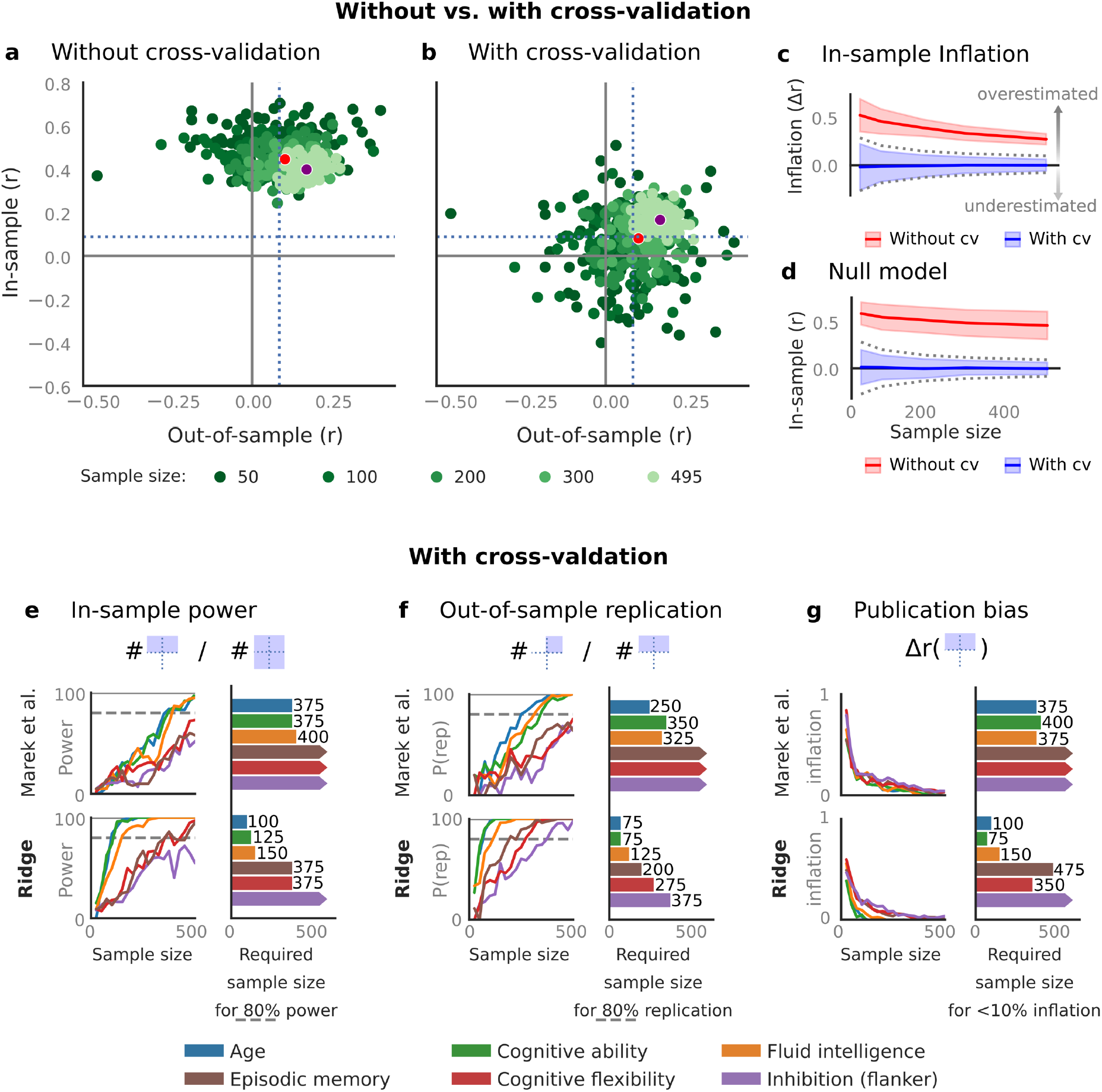
Multivariate BWAS provide unbiased effect sizes and high replicability with low-moderate sample sizes. (**a**) In-sample effects in multivariate BWAS are only inflated if estimates are obtained without cross-validation. **(b)** Cross-validation fully eliminates in-sample effect size inflation and, as a consequence, provides higher replicability. Each point in (a) and (b) corresponds to one bootstrap subsample, as in Fig. 4b of Marek et al. Dotted lines denote the threshold for p=0.05 with n=495. **(c)** The inflation of in-sample effect size obtained without cross-validation (red) is reduced, but does not disappear, at higher sample sizes. Conversely, cross-validated estimates (blue) are slightly pessimistic with low sample sizes and become quickly unbiased as sample size is increased. **(d)** Without cross-validation, in-sample effect size estimates are non-zero (r≈0.5, red) even when predicting permuted outcome data. Cross-validation eliminates systematic bias across all sample sizes (blue). Dashed lines in (c) and (d) denote 95% parametric confidence intervals, and shaded areas denote bootstrap and permutation-based confidence intervals. **(e-f)** Cross-validated analysis reveals that sufficient in-sample power **(e)** and out-of-sample replication probability (P(rep)) **(f)** can be achieved for a variety of phenotypes at low or moderate sample sizes. 80% power and P(rep) are achievable in <500 participants for half the phenotypes tested (colored bars) using the prediction algorithm in Marek et al. (top panels in (e) and (f), sample size required for 80% power or P(rep) shown). Other phenotypes require sample sizes >500 (bars with arrows). Power and P(rep) can be substantially improved with a ridge regression-based model recommended in some comparison studies^10,11^ (bottom panels in (e) and (f)), with 80% power and P(rep) with sample sizes as low as n=100 and n=75, respectively, when predicting cognitive ability, and sample sizes between 75 and 375 for other investigated variables, except inhibition assessed with the flanker task. **(g)** We estimated interactions between sample size and publication bias by computing effect size inflation (r_discovery_ - r_replication_) only for those bootstrap cases where prediction performance was significant (p>0.05) in the replication sample. Our analysis shows that the effect size inflation due to publication bias is modest (<10%) with <500 participants for half the phenotypes using the Marek et al. model and all phenotypes but the flanker using the ridge model.

Given these considerations, how many participants are generally required for multivariate BWAS? The answer to this question depends on the reliability of both phenotypic and brain measures, the size of the effects linking them, the algorithm and model-selection steps employed, and the use cases for the resulting brain measures. For example, multivariate models trained on as few as 20 participants^7^ can have high reliability^8^, broad external validity and large effect sizes in independent samples^9^, when predicting behavioral *states* within-person rather than *traits*. In this case the benefit of large samples is primarily in accurately estimating local brain weights (model parameters)^10^ rather than increasing out-of-sample accuracy. Even when predicting trait-level phenotypes, as Marek et al. did, we estimate that sample sizes of 75-500 are sufficient to achieve high statistical power and replicability (e.g., 80%), and to mitigate effect size inflation due to publication bias, in many cases.

The basis for these estimates is shown in Figure 1e-g. When in-sample estimates were cross-validated, the main multivariate BWAS model used by Marek et al.—principal components-based reduction of bivariate connectivity followed by support vector regression—showed 80% in-sample power and 80% out-of-sample replication probability (P(rep)) at N < 500 for three of six phenotypes we examined (age, cognitive ability, and fluid intelligence). However, this model has been shown to be disadvantageous in some comparison studies^10,11^. Therefore, we performed the same power and sample-size calculations for multivariate BWAS with another approach: ridge regression on partial correlation matrices with a default shrinkage parameter of 1. Though this approach is still likely suboptimal^10,11^ (we avoided testing other models to avoid overfitting), it substantially improved power (Figure 1e, bottom), independent replication probability (Figure 1f, bottom), and resistance to inflation due to publication bias (Figure 1g, bottom). 80% power and P(rep) were achieved at sample sizes from 75 – 150 for age, cognitive ability, and fluid intelligence, and sample sizes <400 for all phenotypes except inhibition measured by the flanker task (a measure based on reaction-time differences with low reliability^12^). In sum, multivariate BWAS are unbiased when accuracy is assessed with appropriate cross-validation procedures and often sufficiently powered with modest sample sizes.

These quantitative differences in required sample size could translate into large, qualitative differences in the types of neuroimaging studies considered viable in future efforts. If >1,000 participants are required for viable BWAS, studies on many topics and populations may never be performed at all, including long-term drug abuse, fibromyalgia, transgender individuals, fragile X syndrome, functional neurological disorders, domain expertise, and more. BWAS based on innovative new tasks, such as those suited for measuring drug craving or learning and generalization rates in computational psychiatry, may be difficult to justify in large-sample studies. Requiring sample sizes larger than necessary could stifle innovation.

We agree with Marek et al. that small-sample studies are important for understanding the brain bases of tasks and mental states^7–9^, and for prototyping new tasks and measures. However, we are substantially more optimistic about the promise of using multivariate BWAS for trait-level behavioral and clinical associations as well^13,14^. Sample sizes of 100-400 are potentially achievable even in moderately rare and vulnerable populations. In addition, several current trends may further increase the viability of small-sample multivariate BWAS, including (a) new phenotypes, (b) feature-learning methods and algorithms with larger effect sizes^11^, (c) models that target within-person variation in symptoms and behavior to improve between-person predictions^2^ and (d) hybrid strategies for improving prediction like meta-matching^15^. All of these can dramatically improve reliability and effect sizes, further increasing the feasibility of BWAS on special populations and in studies using innovative approaches.

## Acknowledgements

We thank Marek et al. for sharing the analysis code and for the thoughtful discussions in relation to our commentary.

## Contributions

Conception and data analysis: T.S.

Manuscript writing, revising: T.S., U.B., T.W.

## Data Availability

Analysis is based on preprocessed data provided by the Human Connectome Project, WU-Minn Consortium (principal investigators: D. Van Essen and K. Ugurbil; 1U54MH091657) funded by the 16 NIH institutes and centers that support the NIH Blueprint for Neuroscience Research; and by the McDonnell Center for Systems Neuroscience at Washington University. All data used in the present study are available for download from the Human Connectome Project (www.humanconnectome.org). Users must agree to data use terms for the HCP before being allowed access to the data and ConnectomeDB; details are provided at https://www.humanconnectome.org/study/hcp-young-adult/data-use-terms.

## Code Availability

All analysis code used in the current study is available at the following github repository (release: v0.3): https://github.com/spisakt/BWAS_comment

