## Supplementary Information for "Replicable multivariate BWAS with moderate sample sizes"

**ARISING FROM**: Marek et al., Reproducible brain-wide association studies require thousands of individuals. https://doi.org/10.1038/s41586-022-04492-9 (2022)

### Supplementary Figures

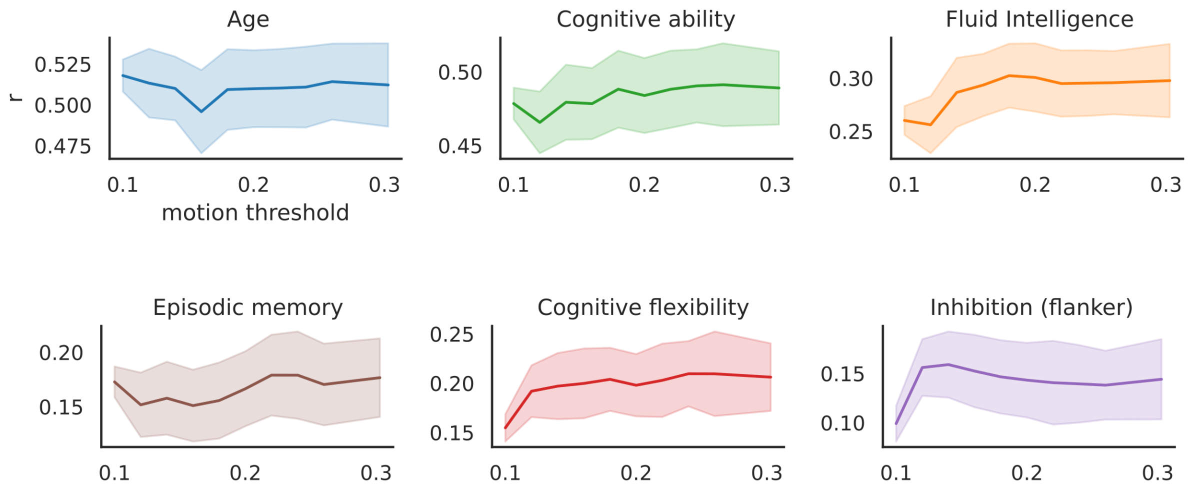


***Supplementary Figure 1.*** *Dependence of out-of-sample predictive performance of the proposed multivariate approach as a function of in-scanner motion (mean root mean squared motion estimates) threshold, for all investigated variables. Sample size is equal for all motion thresholds (N=375 for both the discovery and replication samples; i.e. the number of participants passing the lowest threshold). Analysis was repeated 100 times on random samples (without replacement).*

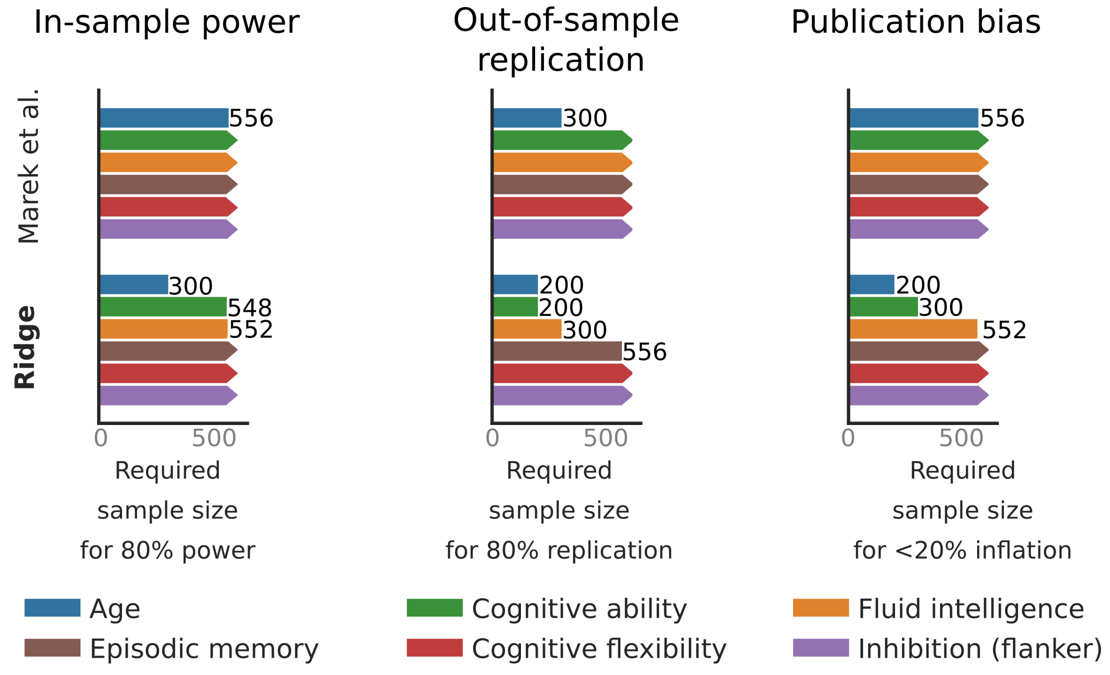


***Supplementary Figure 2.*** *Cross-validated analysis based on cortical thickness measures revealed that sufficient in-sample power (left) and out-of-sample replication probability (P(rep)) (middle) can be achieved for a variety of phenotypes at low or moderate sample sizes. With Ridge regression, 80% power and P(rep) are achievable in <600 participants for age when using the prediction algorithm in Marek et al. (top panels in (e) and (f), sample size required for 80% power or P(rep) shown). Other phenotypes require sample sizes >600 (bars with arrows). Power and P(rep) can be substantially improved with a ridge regression-based model (bottom panels in (e) and (f)), with 80% power and P(rep) with sample sizes as low as n=548 and n=200, respectively, when predicting cognitive ability, and sample sizes between 200 and 556 for other investigated variables, except cognitive flexibility and inhibition assessed with the flanker task. We estimated interactions between sample size and publication bias (right) by computing effect size inflation ($r_{discovery}$ - $r_{replication}$) only for those bootstrap cases where prediction performance was significant (p>0.05) in the replication sample. Our results show that the effect size inflation due to publication bias is modest (<20%) with <500 participants for half the phenotypes using the Ridge model.*

### Supplementary Methods

The Human Connectome Project dataset contains imaging and behavioral data of approximately 1200 healthy subjects[1]. Preprocessed resting state fMRI connectivity data (partial correlation matrices) as published with the HCP1200 release (N=999 participants with functional connectivity data) were used to build models that predict age, cognitive ability, episodic memory fluid intelligence, cognitive flexibility, and inhibition (flanker) scores.

Functional connectivity features were either (i) obtained via full Pearson's correlation coefficient or (ii) via partial correlation, across 100 group-independent component analysis based regions[2].

Cortical thickness was analyzed with the 'Freesurfer'[3], in 63 regions, as defined in the 68 regions of the Desikan-Killiany Atlas[4].

No participants were excluded due to high in-scanner motion. This decision was based on a bootstrap analysis of the dependence of out-of-sample predictive performance on the threshold for excluding participants. (Supplementary Figure 1).

Discovery and replication samples with various sample sizes were randomly sampled form all available participants 100 times for each sample size (ranging from 25 to ~500). On the discovery sample two machine learning models were evaluated via cross validation. The first model consisted of a principal component analysis retaining 50% of the total feature variance and a support vector regression and was trained on cortical thickness values and the full Pearson correlation features (as in Marek et al.). The second model was a Ridge regression (with shrinkage parameter set to 1), trained on cortical thickness values (Supplementary Figure 2) and functional connectivity features.

Model performance in the discovery sample was evaluated by averaging the correlation coefficient between the predicted and observed values in the test set, across all folds.

The models were then fit on the whole discovery sample and used to predict the replication sample.
